# A functional drug re-purposing screening identifies carfilzomib as a drug preventing 17β-estradiol: ERα signaling and cell proliferation in breast cancer cells

**DOI:** 10.1101/133256

**Authors:** Claudia Busonero, Stefano Leone, Cinzia Klemm, Filippo Acconcia

## Abstract

Most cases of breast cancer (BC) are estrogen receptor α-positive (ERα+) at diagnosis. The presence of ERα drives the therapeutic approach for this disease, which often consists of endocrine therapy (ET). 4OH-Tamoxifen and faslodex (*i.e.,* fulvestrant - ICI182,780) are two ETs that render tumor cells insensitive to 17β-estradiol (E2)-dependent proliferative stimuli and prevent BC progression. However, ET has limitations and serious failures in different tissues and organs. Thus, there is an urgent need to identify novel drugs to fight BC in the clinic. Re-positioning of old drugs for new clinical purposes is an attractive alternative for drug discovery. For this analysis, we focused on the modulation of intracellular ERα levels in BC cells as target for the screening of about 900 Food and Drug Administration (FDA) approved compounds that would hinder E2:ERα signaling and inhibit BC cell proliferation. We found that carfilzomib induces ERα degradation and prevents E2 signaling and cell proliferation in two ERα+ BC cell lines. Remarkably, the analysis of carfilzomib effects on a cell model system with an acquired resistance to 4OH-tamoxifen revealed that this drug has an antiproliferative effect superior to faslodex in BC cells. Therefore, our results identify carfilzomib as a drug preventing E2:ERα signaling and cell proliferation in BC cells and suggest its possible re-position for the treatment of ERα+ BC as well as for those diseases that have acquired resistance to 4OH-tamoxifen.

## Introduction

17β-estradiol (E2) regulates the physiology of reproductive and non-reproductive organs both in males and in females by binding to the estrogen receptors (*i.e.,* ERα and ERβ) and thus it is a general regulator of body homeostasis [1]. The E2 pleiotropic nature resides in its ability to elicit diverse intracellular effects (including cell proliferation) through multifaceted signaling mechanisms. As nuclear receptors, ERα and ERβ localize into the nucleus where they mediate changes in gene transcription profiles in response to E2 (*i.e.,* nuclear effects). ERα and ERβ also localize at the cell plasma membrane where they activate E2 extra-nuclear signals. Nuclear and extra-nuclear events cross-talk and are both required for cell and organ physiology [1-3]. Many E2 physiological effects occur while E2 finely tunes ERα and ERβ intracellular levels. This modulation is specific for each receptor subtype with E2 reducing ERα intracellular levels and inducing ERβ intracellular accumulation. Therefore, E2 signaling and its resulting physiological functions further depend on the dynamic temporal variations in ERs cell content [1, 4, 5]. In turn, deregulation of both E2 intracellular signaling and control of ERs intracellular levels contributes to the development of diverse endocrine-related diseases including breast cancer (BC) [1-5].

In 70% of all cases, BC growth depends on E2 signaling through ERα (*i.e.,* ERα+ tumors) as E2 is a mitogen for BC cells, works as a survival and anti-apoptotic factor and induces cell invasion and migration [6]. The clinical approach for ERα+ tumors targets different aspects of oncogenic E2:ERα signaling: aromatase inhibitors (AIs) are used to inhibit E2 synthesis thus reducing systemic E2 availability; 4OH-tamoxifen (*i.e.,* the actual mainstay for treatment of ERα+ tumors) is the prototype of the selective ER modulators (SERMs), it binds ERα and inhibits receptor to work as a transcription factor, thus blocking E2-dependent gene expression; and faslodex (*i.e.,* fulvestrant or ICI182,780) is the prototype of the selective ER down-regulators (SERDs), physically engages with ERα and induces its 26S proteasome-dependent degradation [7, 8], thus eliminating ERα from BC cells.

However, the use of such drugs has limitations: AIs are typically reserved for postmenopausal women with BC and produce musculoskeletal failures, and 4OH-tamoxifen and faslodex possess serious side effects (*e.g.,* endometrial cancer for 4OH-tamoxifen) and often determine the development of BC cells resistant to such anti-cancer ET [9, 10]. Consequently, considerable drug discovery research efforts aimed to identify novel SERMs and SERDs but 4OH-tamoxifen and faslodex stand still as the unique ERα ligands available for BC treatment [7, 8]. Thus, additional drugs to treat BC are required. Novel compounds that inhibit ERα+ BC progression can be identified either by searching for natural or synthetic compounds that bind to and/or inhibit ERα or by defining and targeting a critical parameter required for E2:ERα signaling-induced cell proliferation.

We previously hypothesized that such parameter can be the modulation of ERα intracellular levels [1, 11] as i) it is known since many years that both artificial ERα down-modulation in ERα+ cell lines and its over-expression in ERα-negative cells cause E2 insensitivity or cell death [12, 13]; ii) ERα ligands influence receptor intracellular concentration: E2 reduces ERα intracellular levels and fuels cell proliferation [4, 14, 15], 4OH-tamoxifen and faslodex alter ERα content, with the former stabilizing it, the latter de-stabilizing it and either drugs blocking E2-induced BC cell proliferation [4]; iii) molecules (*e.g.,* chloroquine) that do not bind to ERα influence ERα levels by affecting specific cellular pathways and prevent E2-induced BC cell proliferation [15, 16]; and iv) reduction in the intracellular content of proteins with functions unrelated to E2 signaling (*e.g.,* endocytic proteins) can change ERα content and inhibit E2-induced BC cell proliferation [17, 18]. In turn, all this evidence suggests that molecules that directly or indirectly deregulate the control mechanisms for ERα intracellular abundance could have in principle the potential to inhibit E2-dependent proliferation in BC cells.

Herein, we challenged this hypothesis by screening a compound library with the aim to identify molecules that modify ERα intracellular levels. We report the discovery of carfilzomib (Kyprolis®) as an ERα-degrading drug that prevents the basal and E2-induced BC cell proliferation.

## Materials and Methods

### Cell culture and reagents

17β-estradiol, DMEM (with and without phenol red) and fetal calf serum were purchased from Sigma-Aldrich (St. Louis, MO). Bradford protein assay kit as well as anti-mouse and anti-rabbit secondary antibodies were obtained from Bio-Rad (Hercules, CA). Antibodies against ERα (F-10 mouse), cyclin D1 (H-295 rabbit), cathepsin D (H75 rabbit), pS2 (FL-84 rabbit), were obtained from Santa Cruz Biotechnology (Santa Cruz, CA); anti-vinculin antibody was from Sigma-Aldrich (St. Louis, MO). Chemiluminescence reagent for Western blotting was obtained from BioRad Laboratories (Hercules, CA, USA). Faslodex (*i.e.,* fulvestrant or ICI182,780) and 4OH-tamoxifen were purchased by Tocris (USA). FDA-approved drug library was purchased by Selleck Chemicals (USA). All the other products were from Sigma-Aldrich. Analytical- or reagent-grade products were used without further purification. The identities of all of the used cell lines [*i.e.,* human breast carcinoma cells (MCF-7; ZR-75-1)] were verified by STR analysis (BMR Genomics, Italy).

### Cellular and biochemical assays

Cells were grown in 1% charcoal-stripped fetal calf serum medium for 24 hours and then stimulated with E2 at the indicated time points. Where indicated, cells were treated with E2 (1 nM), carfilzomib (100 nM) or Faslodex (*i.e.,* fulvestrant or ICI182,780) (ICI) (100 nM). Protein extraction and biochemical assays were performed as described previously [11]. For cell number analysis, the CyQUANT® Cell Proliferation Assay (Life Technologies) was used according to the manufacturer’s instructions. Specifically, MCF-7, ZR-75-1 and 4OH-Tamoxifen resistant MCF-7 cells (Tam-Res) were treated as described below. After counting 3,000 cells were plated in 96-well plates in triplicate. After 4 hours, the CyQUANT® Assay was performed at time 0 (*i.e.,* plated cells). In other 96-well plates, drugs were administered to cells in 1% charcoal-stripped fetal calf serum medium, and the CyQUANT® assay was performed after 48 hour. For growth curves analysis, parental and Tam-Res MCF-7 cells were plated in growing medium and counted at the indicated time points after drug administration. Western blotting analyses were performed by loading 20-30 μg of protein on SDS-gels. Gels were run and transferred to nitrocellulose membranes with Biorad Turbo-Blot semidry transfer apparatus. Immunoblotting was carried out by incubating membranes with 5% milk (60 min), followed by incubation o.n. with the indicated antibodies. Secondary antibody incubation was continued for an additional 60 min. Bands were detected using a Biorad Chemidoc apparatus.

### Cell cycle analysis

After treatments, cells were grown in 1% FBS for 24 hours, harvested with trypsin, and counted to obtain 106 cells per condition. Then, the cells were centrifuged at 1500 rpm for 5 min at 4°C, fixed with 1 ml ice-cold 70% ethanol and subsequently stained with PI buffer (500 μg/ml Propidium Iodide, 320μg/ml RNaseA, in 0.1% Triton X in PBS). DNA fluorescence was measured using a CytoFlex flow cytometer and the cell cycle analysis was performed by CytExpert v1.2 software (Beckman Coulter).

### Statistical analysis

A statistical analysis was performed using the ANOVA (One-way analysis of variance and Tukey’s as post-test) test with the InStat version 3 software system (GraphPad Software Inc., San Diego, CA). Densitometric analyses were performed using the freeware software Image J by quantifying the band intensity of the protein of interest respect to the relative loading control band (*i.e.,* vinculin or tubulin) intensity. In all analyses, *p* values < 0.01 were considered significant, except of the densitometric analyses with a choosen threshold of p < 0.05. Z’ factor and robust Z scores calculation was performed according to [19] and [20], respectively.

## Results

### FDA-approved drug library screen

In order to test if the modulation of ERα levels could be used as a pharmacological target for identifying anti-ERα+ BC drugs [1, 11], we applied a library of Food and Drug Administration (FDA)-approved drugs to ERα expressing ductal carcinoma cells (MCF-7) (Fig. 1A). MCF-7 cells were 1 hour pre-treated with each drug (100 nM) before E2 (1 nM) administration and further grown for 24 hours (Fig. 1B). Subsequently, both intracellular ERα and cathepsin D levels were evaluated by Western blotting to contemporarily test the effect of the compounds both on ERα levels and on the levels of an ERα transcriptional target [21]. The library was composed of 1018 compounds organized into different pharmacological categories (Fig. 1A). We excluded from this array those molecules that are intended for cancer treatment as well as 4OH-tamoxifen, faslodex and aromatase inhibitors and screened about 900 drugs. Overall, we performed 162 Western blots and densitometrically quantitated ERα levels using vinculin as loading control. After quantitation, we set as 100% the control value for ERα levels in DMSO-treated (CTR) MCF-7 cells in each Western blot. As expected [4], E2 reduced ERα content in MCF-7 cells to 24,09% ± 3,94% with respect to control and analysis of controls and E2 treated samples resulted in a Z’ factor equal to 0.84, which identifies a large separation between the test groups [19].

**Figure 1.**
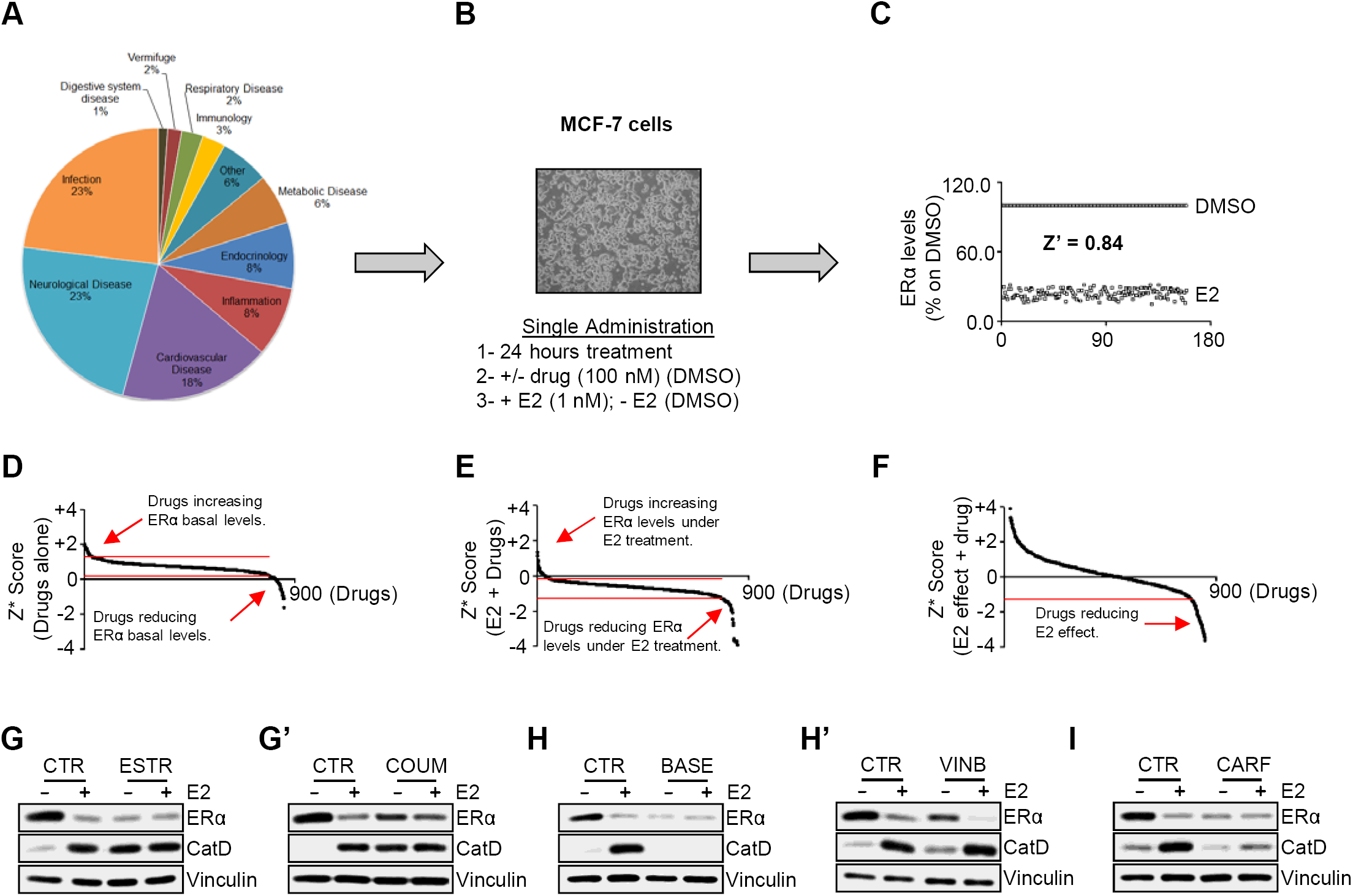
The impact of FDA-approved drugs on ERα levels in MCF-7 cells. (A) Pie diagrams depicting the pharmaceutical categories in the library. (B) Synthetic protocol used for drug administration to MCF-7 cells. (C) Assay data depicting the quantitation of ERα levels in each one of the 162 performed Western blots. Z’ is the Z factor for negative (DMSO) and positive (E2) controls in each Western blot analysis. Robust Z scores (Z*) graphs for drug-treated samples alone (D), in combination with E2 (E) or for the effect of E2 within each drug-treated sample (*i.e.,* E2+drug / drug alone) (F); red arrows indicate the Z* for drugs considered as positive hits. Red lines indicate the threshold used for analysis. Western blot analyses of ERα and cathepsin D (Cat D) expression levels in MCF-7 cells treated for 24 hours with E2 (1 nM) both in the presence or in the absence of 100 nM estriol (ESTR) (G), coumarin (COUM) (G’), basedoxifene (BASE) (H), vinblastine (VINB) (H’) and carfilzomib (CARF) (I) or vehicle (DMSO-CTR). The loading control was done by evaluating vinculin expression in the same filter.

We next calculated robust Z score [20] for each sample (data not shown) and then separated the robust Z scores for drug-treated samples alone or in combination with E2 (Fig. 1D and Fig. 1E). In addition, robust Z scores for the effect of E2 within each drug-treated sample (*i.e.,* E2+drug / drug alone) were also calculated (Fig. 1F). Thresholds as indicated in figure 1D, 1E and 1F were set to identify groups of drugs that either reduced or increased ERα intracellular levels both in the basal and in E2-treated samples as well as to identify those drugs that reduced the effect of E2 in the presence of each drug. By using these limits, we shortlisted 53 drugs that differentially modulated ERα intracellular levels. Eight compounds were further excluded as they have estrogenic activity (*i.e.,* estriol, coumarin [22]) (Fig. 1G and Fig. 1G’), are known ERα down-regulators (*i.e.,* basedoxifene [23] and vinblastine [24]) (Fig. 1H and Fig. 1H’) or belong to a pharmaceutical category with scarce indication for BC treatment (Table 1) (*i.e.,* progestins: norethindrone and megestrol; immunosuppressants: bethametasone and mycophenolate).

Next, the resulting 45 drugs were tested for their ability to affect cell proliferation. The number of MCF-7 cells was detected after 48 hours of E2 (1 nM) administration both in the presence and in the absence of the treatment with each one of the 45 selected drugs (100 nM). Seven compounds significantly (p<0.05) inhibited basal and/or E2-induced cell proliferation. Among them, we found emetine [25], and two compounds with therapeutic indications for cardiovascular diseases. The remaining three drugs are used for treating respiratory, neurological and infectious pathologies.

Interestingly, we also identified carfilzomib (Kyprolis®) (Fig. 1I) as a drug inducing ERα degradation and preventing E2-induced BC cell proliferation. Thus, we decided to focus on this second generation 26S proteasome inhibitor, as its use is approved to treat blood tumors [26] but its effects on E2:ERα signaling in BC cells are unknown.

### Carfilzomib changes ERα levels and perturbs E2-induced proliferation in MCF-7 and ZR-75-1 cells

Initial experiments were performed to validate the ability of carfilzomib to reduce ERα intracellular levels in BC cells. Both MCF-7 and ZR-75-1 cells were treated for 1 hour with carfilzomib before 24 hours E2 administration. Faslodex (*i.e.,* ICI182,780 - ICI) was also introduced in the experiments as a control for ERα degradation. Figure 2A, 2B and 2C show that E2, ICI and carfilzomib reduced ERα intracellular levels in each cell line. Remarkably, while ICI inhibited E2-dependent ERα degradation, carfilzomib administration did not affect the ability of E2 to reduce ERα intracellular content (Fig. 2A, 2B and 2C). As expected by a 26S proteasome inhibitor [26], carfilzomib administration to MCF-7 and ZR-75-1 cells increased the total amount of total ubiquitinated species (Fig. 2A and 2B).

**Figure 2.**
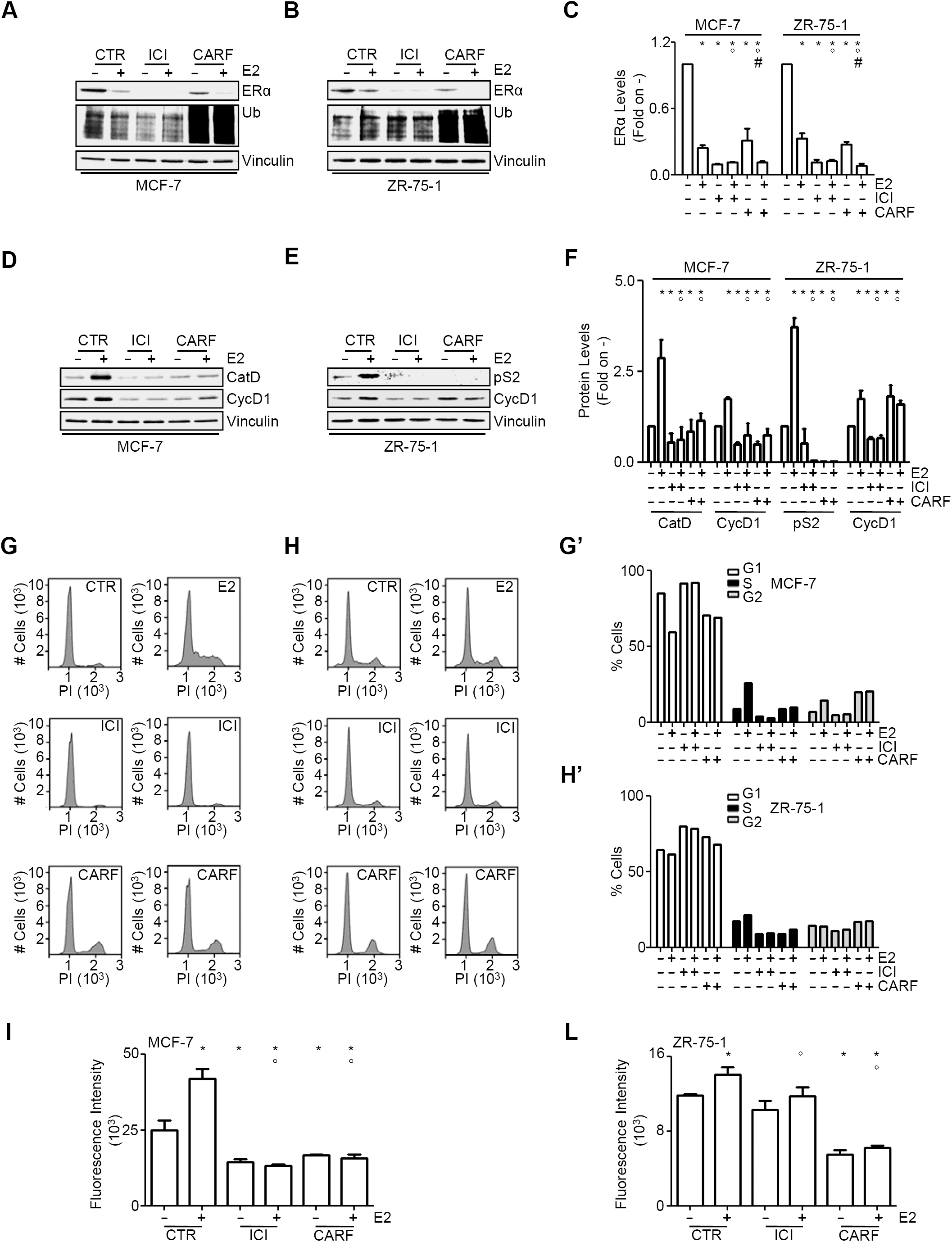
The effect of carfilzomib on ERα degradation and E2-induced cell proliferation in MCF-7 and ZR-75-1 cells. Western blotting and relative densitometric analyses of ERα, cathepsin D (Cat D), presenelin 2 (pS2), cyclin D1 (Cyc D1) and ubiquitin (Ub) expression levels in starved MCF-7 (A, D, C, F) and ZR-75-1 (B, E, C, F) cells treated for 24 hours with E2 (1 nM) both in the presence or in the absence of either ICI182,780 (ICI – 100 nM), carfilzomib (CARF – 100 nM) or vehicle (DMSO-CTR). The loading control was done by evaluating vinculin expression in the same filter. Cell cycle (G, G’, H, H’) and cell number (I, L) analyses in MCF-7 and ZR-75-1 cells, respectively, treated as described in (A-F). * indicates significant differences with respect to the CTR sample; ° indicates significant differences with respect to the E2 sample; # indicates significant differences with respect to the CARF sample. All experiments were performed in triplicates. Data are the mean ± standard deviations with a p value < 0.01.

Next, we tested if the carfilzomib-mediated reduction in ERα levels could correlate to a reduction in the ability of the cells to functionally respond to E2 stimulation. Therefore, the expression of two well-known estrogen response element (ERE)-containing genes [*i.e.,* presenelin2/TIFF (pS2) and cathepsin D (Cat D)] was tested under carfilzomib treatment. As shown in figure 2D, 2E and 2F, the E2-induced accumulation of cathepsin D and pS2 was significantly reduced after both ICI and carfilzomib treatment in MCF-7 and ZR-75-1 cells, respectively. ERα transcriptional activity also occurs on non-ERE-containing genes. This indirect mode of regulation is mediated by ERα binding to other transcription factors (*e.g.,* AP-1 and Sp-1) that are activated by E2-induced extra-nuclear signaling. Thus, MCF-7 and ZR-75-1 cells were used to study the effect of carfilzomib on the E2-dependent control of cyclin D1 intracellular levels, a recognized E2 target gene, which promoter does not contain any ERE sequence [27, 28]. Figure 2D, 2E and 2F show that the E2-dependent induction of cyclin D1 protein levels was reduced in ICI- and carfilzomib-treated cells. Interestingly, basal cyclin D1 expression was increased in ZR-75-1 cells but not in MCF-7 cells, suggesting a different sensitivity of BC cell lines to carfilzomib (Fig. 2D, 2E and 2F). Therefore, carfilzomib-induced reduction in ERα levels impacts on E2-induced receptor transcriptional activity.

As E2-dependent transcription of cyclin D1 is necessary for hormone-dependent induction of cell cycle progression [15, 27-29], the ability of E2 to induce cellular proliferation in MCF-7 and ZR-75-1 cells was further evaluated in the presence of carfilzomib. As expected, cell cycle analysis revealed that ICI and carfilzomib administration to MCF-7 and ZR-75-1 cells prevents E2-triggered cell cycle progression. Remarkably, carfilzomib increased the percentage of cells in the G2 phase of the cell cycle irrespective of E2 administration while ICI determined an accumulation of cells in the G1 phase of the cell cycle (Fig. 2G, 2H, 2G’ and 2H’). Accordingly, the E2-depedent increase in cell number was prevented by both ICI and carfilzomib administration in either BC cell lines. Notably, carfilzomib administration partially reduced the basal cell number in MCF-7 and ZR-75-1 cells (Fig. 2I and 2L) while ICI only affected basal MCF-7 cell number.

Taken together, these data demonstrate that carfilzomib administration to BC cells interferes with the ability of E2 to induce cell proliferation.

### Carfilzomib changes ERα levels and prevents cell proliferation in 4OH-tamoxifen-resistant MCF-7 cells

Because carfilzomib inhibits E2-induced ERα-mediated cell proliferation in BC cells, we next sought to determine the effect of this drug on 4OH-tamoxifen-resistant (Tam-Res) MCF-7 cells. The growth of Tam-Res MCF-7 cells, which were previously selected by continuous administration of 100 nM 4OH-tamoxifen [30], was analyzed under different doses of 4OH-tamoxifen treatment for 72 hours in comparison to parental MCF-7 cells. Figure 3A shows that while 4OH-tamoxifen significantly inhibited the growth of parental MCF-7 cells it did not produce any effect on the number of Tam-Res MCF-7 cells at any of the 4OH-tamoxifen concentrations used. Additionally, as it has been previously shown that a reduction in B cell lymphoma 2 (Bcl-2) levels is observed in many MCF-7 cells selected to be resistant to 4OH-tamoxifen [31-33], we next measured the levels of Bcl-2 in parental and Tam-Res MCF-7 cells. As expected, Bcl-2 levels were undetectable in Tam-Res MCF-7 cells (Fig. 3A, inset). These data confirm that the Tam-Res MCF-7 cells display some of the typical features of the MCF-7 cells that acquired resistance to 4OH-tamoxifen *in vitro*.

**Figure 3.**
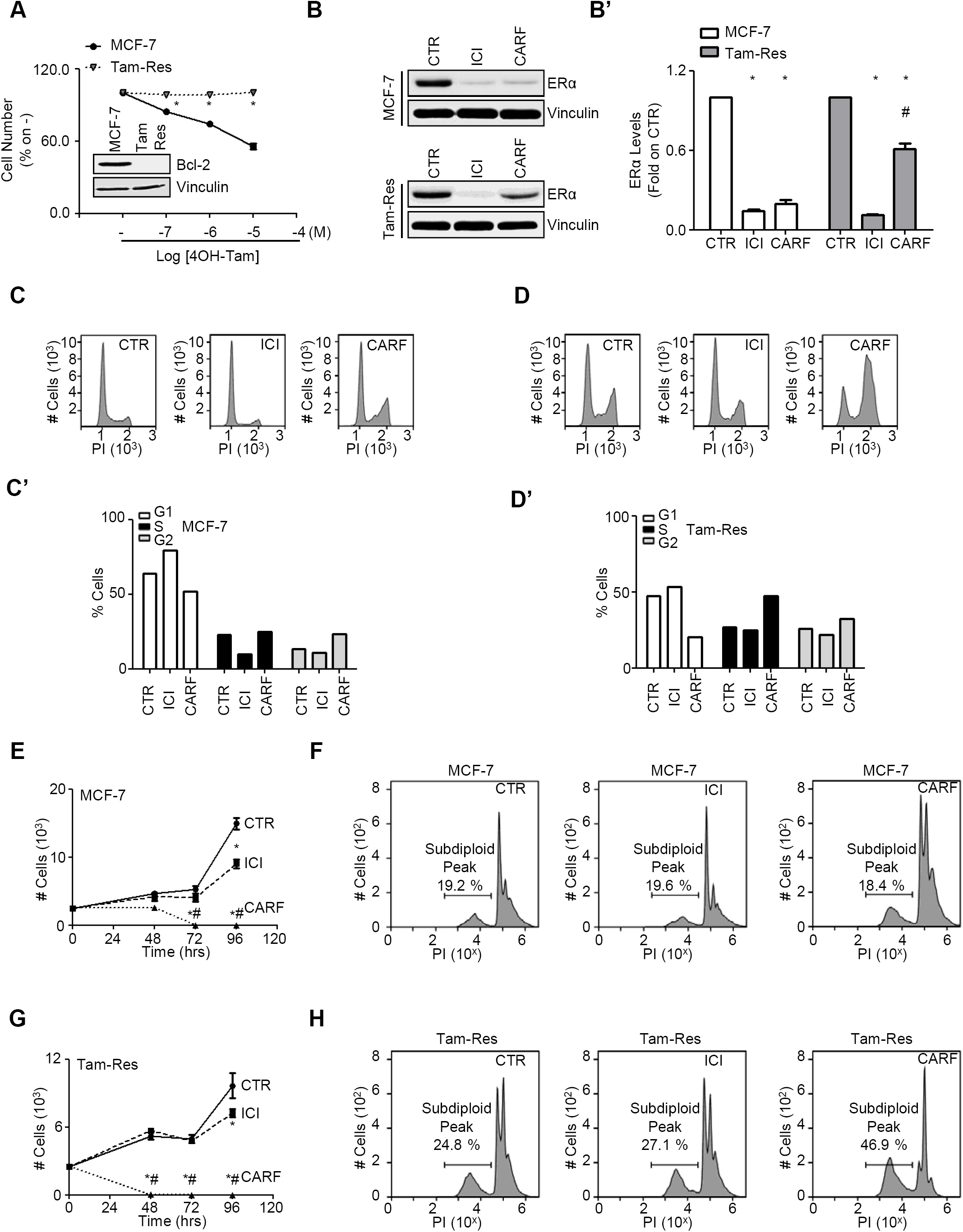
The effect of carfilzomib on ERα degradation and cell proliferation in parental and 4OH-tamoxifen-resistant MCF-7 cells. (A) Cell number analyses in parental (MCF-7) 4OH-tamoxifen resistant (Tam-Res) MCF-7 cells after 72 hours treatment with the indicated doses of 4OH-tamoxifen (4OH-Tam). (A-inset) Western blot analyses of Bcl-2 levels in parental (MCF-7) 4OH-tamoxifen resistant (Tam-Res) MCF-7 cells. (B) Western blotting and relative densitometric analyses of ERα expression levels in parental (MCF-7) 4OH-tamoxifen resistant (Tam-Res) MCF-7 cells treated for 24 hours in complete medium with either ICI182,780 (ICI – 100 nM), carfilzomib (CARF – 100 nM) or vehicle (DMSO-CTR). The loading control was done by evaluating vinculin expression in the same filter. Cell cycle (C, C’, D, D’), growth curves (E, G) and sub-G1 (F, H) analyses in parental (MCF-7) 4OH-tamoxifen resistant (Tam-Res) MCF-7 cells, respectively, treated with either ICI182,780 (ICI – 100 nM), carfilzomib (CARF – 100 nM) or vehicle (DMSO-CTR). * indicates significant differences with respect to the CTR sample; # indicates significant differences with respect to the CARF sample. All experiments have been repeated three times. Data are the mean ± standard deviations with a p value < 0.01.

Next, the levels of ERα were measured after 24 hours treatment with carfilzomib in comparison with ICI. Figure 3B and 3B’ shows that both ICI and carfilzomib reduced ERα intracellular levels in parental MCF-7 cells to the same extent while ICI was more effective than carfilzomib in reducing ERα content in Tam-Res MCF-7 cells. Moreover, the effect of carfilzomib in reducing receptor concentration was less pronounced in Tam-Res MCF-7 cells (ERα levels = 61.26 % ± 0.05) than that observed in parental MCF-7 cells (ERα levels = 19.67 % ± 0.03). These data demonstrate that carfilzomib administration determine a reduction in ERα levels also in Tam-Res MCF-7 cells and further suggest that the ERα expressed in Tam-Res MCF-7 cells is more resistant to carfilzomib-induced degradation.

Cell cycle analysis was next performed to study the impact of carfilzomib on the cell cycle in Tam-Res MCF-7 cells. As expected ICI administration determined an increase in the percentage of both parental and Tam-Res MCF-7 cells in the G1 phase of the cell cycle. On the contrary, carfilzomib increased the number of cells in the S-G2 transition phase of the cell cycle in either cell lines (Fig. 3C, 3D, 3C’ and 3D’). These data suggest that carfilzomib could prevent the proliferation of both parental and Tam-Res MCF-7 cells. Therefore, we next performed growth curves analysis in either cell lines in the presence or in the absence of either ICI or carfilzomib. Figure 3E and 3G show that while ICI significantly reduced the proliferation of each cell line after 96 hours of compound administration, treatment with carfilzomib dramatically reduced the number of both parental and Tam-Res MCF-7 cells. Interestingly, within the first 48 hours of treatment, carfilzomib prevented the growth of parental MCF-7 cells while it reduced the number of Tam-Res MCF-7 cells. These data indicate that carfilzomib inhibits the growth of both parental and Tam-Res MCF-7 cells with Tam-Res MCF-7 cells being more sensitive than the parental counterparts. Accordingly, 24 hours administration of carfilzomib but not of ICI significantly increased the subdiploid peak only in the cell cycle of Tam-Res MCF-7 cells (Fig. 3F and 3H).

Overall, these data indicate that carfilzomib induces the death of parental and Tam-Res MCF-7 cells and further suggest that this drug works better than ICI in preventing BC cell proliferation.

## Discussion

E2 controls several physiological functions, among which cellular proliferation through its binding to ERα and to ERβ. E2:ERβ complex drives anti-proliferative effects, while E2:ERα complex induces proliferative and anti-apoptotic effects. E2-induced cell proliferation is accompanied by the hormone-dependent ERα degradation. These effects are controlled by nuclear and extra-nuclear ERα signaling, which depends on nuclear and plasma membrane ERα localization. Regulation of ERα intracellular levels occurs as the result of a vast array of molecular processes that include either 26S proteasome-based, lysosome-based and autophagy-based pathways, genetic or epigenetic mechanisms. Notably, deregulation of the mechanisms controlling ERα levels is an emerging feature of endocrine-related cancers in general and of BC in particular [1-5]. Thus, the control of ERα intracellular content is intrinsically connected with the appearance of the E2 effects.

Remarkably, because of its physiological importance, this parameter represents for BC cells such an intrinsic weakness that any factor causing the de-regulation of ERα levels can potentially inhibit E2-dependent proliferation. In fact, the artificial manipulation of ERα content (*i.e.,* over-expression in receptor-devoid cell lines or ERα gene silencing in ERα-expressing cells) induces cell death. Some ERα ligands (*e.g.,* 4OH-tamoxifen and faslodex) change ERα levels and block E2-induced cell proliferation. Recently, we and others have also shown that the interference with pathways not related to ERα obtained by specific molecules (*e.g.,* chloroquine) or by selective gene knock-down (*e.g.,* siRNA against clathrin, caveolins, and dynamin II) modifies ERα intracellular content and prevents E2 proliferative effects [1, 4, 11-18]. On this basis, we have proposed [1, 11, 25] that the modulation of ERα intracellular concentration in BC cells could be used as a pharmacological target to identify anti-tumor drugs.

Here, we challenged this assumption and applied a library of about 900 compounds (FDA-approved drugs) already used in clinical practice to MCF-7 cells by analyzing the ERα intracellular levels. Many investigators have performed high-throughput screenings in BC cells aimed to identify compounds with anti-cancer activity or pathways for drug discovery by using cell proliferation as the readout [34, 35]. To our knowledge, this is the first screen that has exploited the modulation of ERα levels to fish out molecules that change ERα intracellular content. It is important to note here that the entire screen has been based on Western blotting to detect changes in ERα intracellular levels because we assumed this technique as the best and most effective way of getting definitive information on potential drug-induced variations in ERα content. However, Western blotting lacks high throughput potential. Thus, to challenge this parameter with larger compound and/or siRNA libraries, a method to detect ERα levels in (at least) 96-well plate format is desirable. In this respect, we are investigating the possibility to measure ERα levels by in-cell Western blotting [36].

Nonetheless, we have identified 53 drugs changing receptor concentration in MCF-7 cells. Among these drugs, we found basedoxifene and vinblastine that were previously known to induce ERα degradation [23, 24]. These results together with the identification of estriol and coumarin that have known estrogenic effects [22] and modulate ERα intracellular content in MCF-7 cells strongly support the notion that the modulation of ERα levels can be used to find molecules affecting E2:ERα signaling. Next, we searched those compounds that, in addition to change receptor content, were also able to prevent E2-induced BC cell proliferation and shortlisted 7 drugs. Emetine was one of these molecules. Emetine is an alkaloid produced by ipecac roots and used as an anti-parasitic and contraception molecule [37, 38]. We have previously reported that emetine reduces ERα intracellular levels and prevents E2-induced cell proliferation without directly binding to ERα. Therefore, we have concluded that that drugs not necessarily binding to ERα can block E2:ERα signaling to cell proliferation [25].

Another compound identified in the screen that reduces ERα levels and E2-induced BC cell proliferation is carfilzomib (Kyprolis®), an irreversible 26S proteasome inhibitor [26]. Indeed, treatment of two different BC cell lines (*i.e.,* MCF-7 and ZR-75-1 cells) with carfilzomib results in a significant reduction in ERα content and in a consequent block in the ability of E2 to induce ERE and non-ERE containing gene expression (*i.e.,* pS2, cathepsin D and cyclin D1) as well as cell proliferation. On this basis, we conclude that carfilzomib perturbs E2:ERα signaling and dampens hormone-induced cell proliferation in BC cells. Remarkably, the fact that E2 is still able to trigger ERα degradation in the presence of carfilzomib suggests that this drug does not bind to ERα and that its effects depend on the ability of the drug to inhibit the 26S proteasome. Accordingly, treatment of BC cell lines with other inhibitors of 26S proteasome activity (*e.g.,* bortezomib – Velcade®) reduces ERα levels and consequently prevents receptor-dependent signaling [39]. The apparent paradox that 26S proteasome inhibitors reduce the levels of ERα, which turnover is, at least in part, controlled by the 26S proteasome itself can be reconciled by considering that the irreversible inhibition of 26S proteasome leads to a high accumulation of ubiquitinated species, which cells remove by activating autophagy [40, 41]. In this respect, ERα basal turnover is under the control of the autophagic flux [11] and both carfilzomib and bortezomib can activate autophagy [42, 43].

Carfilzomib also works in reducing ERα intracellular levels and cell proliferation in a MCF-7 cell line that has acquired resistance to 4OH-tamoxifen (Tam-Res) [30]. Interestingly, we found that while in Tam-Res MCF-7 cells carfilzomib induces a reduction in receptor levels that is less pronounced than that observed in parental MCF-7 cells, this drug determines a more dramatic effect on the proliferation of Tam-Res MCF-7 cells with respect to the parental ones. In particular, carfilzomib treatment perturbs the cell cycle mainly in the G2 phase and induces an accumulation of fragmented DNA, a canonical sign of apoptosis. Moreover, growth curve analysis revealed that this drug reduces the number of both parental and Tam-Res MCF-7 cells within 48 hours treatment. Therefore, we conclude that carfilzomib kills parental and Tam-Res BC cells.

Another interesting observation reported here is the fact that carfilzomib has an anti-proliferative effect superior to faslodex (*i.e.,* fulvestrant or ICI182,780) in all the BC cell lines used in this study including the Tam-Res cell line. 4OH-tamoxifen is the current mainstay for the treatment of ERα expressing breast tumors. However, at least one-third of women treated with 4OH-tamoxifen for 5 years will relapse within 15 years and the resulting tumors will be insensitive to 4OH-tamoxifen treatment. Faslodex is the only second line ET that is administered to women with a Tam-Res BC [9, 10]. Therefore, our findings suggest that carfilzomib could be useful also for the treatment of patients with ERα+ BC that acquired resistance to 4OH-tamoxifen during antitumor therapy.

In conclusion, we demonstrate here that the selective modulation of ERα intracellular levels can be used as a pharmacological target to identify novel molecules (and/or lead compounds) targeting E2:ERα signaling. Such procedure can be potentially valuable (and cost-effective) to extend the repertoire of drugs for ERα+ BC treatment. As an example of this unconventional approach, present results provide support for further clinical research to evaluate the possibility to re-position carfilzomib as a drug for treating breast cancer including 4OH-tamoxifen-resistant tumors.

## Acknowledgements

The 4OH-Tamoxifen resistant MCF-7 cells (Tam-Res) were a generous gift of Dr Carol Dutkowski, University of Cardiff, England. This study was supported by grants from AIRC-Associazione Italiana Ricerca sul Cancro (MFAG12756) to F.A. and Ateneo Roma Tre to F.A.

